# VariantFlow: a selective-execution engine for efficient population genomic computation on large variant datasets

**DOI:** 10.64898/2026.09.01.748643

**Authors:** Ehsan Estaji, Jian-Feng Mao

## Abstract

Population-scale sequencing now produces variant call sets with thousands of samples and millions of sites, making post-calling analysis a recurring bottleneck. Because the Variant Call Format stores every field of every record together, a tool answering a field-limited question still parses the unused annotations, FORMAT blocks, and per-sample values, which costs time without changing the result. We present VariantFlow, a command-line engine built on selective execution, decoding only the fields the current filter, statistic, or projection requires and preserving original records where possible. On correctness-matched benchmarks on the 1000 Genomes 3 202-sample high-coverage dataset, VariantFlow computed whole-chromosome missingness 8.0–8.5x faster than VCFtools at a constant 8.6 MB of memory, the chromosome 1 summary taking 128 s against 1031 s. An index-assisted `FILTER = PASS` predicate ran 123–273x faster than bcftools where block metadata excludes most of the file. Further gains, reported in Table 1, cover FORMAT-rich filtering on record subsets, linkage disequilibrium, and export-once Parquet queries served through DuckDB. Core population-genetic outputs were byte-identical to VCFtools, per-individual missingness and the site-frequency spectrum matched scikit-allel exactly, and the missing-data-aware *π* and *d*_*XY*_ estimators reproduced pixy’s pairwise counts in every window. By decoding only the fields each operation requires, VariantFlow brings large-cohort post-calling analysis within reach of exploratory work on a single workstation. It complements rather than replaces bcftools, HTSlib, GATK, and VCFtools, and should be of value wherever large cohort VCFs are repeatedly summarised, filtered, or exported. VariantFlow is open source (Rust, MIT OR Apache-2.0) at https://github.com/ehsanestaji/VariantFlow; version 1.5.0 is archived at doi:10.5281/zenodo.21198172.

## 1 Introduction

The volume of genomic variant data is growing rapidly. Population-scale sequencing projects now routinely produce Variant Call Format (VCF) files that describe thousands of samples and millions of variant sites per chromosome (Danecek et al., 2011). Once variants have been called, these files are scanned repeatedly for quality filtering, cohort subsetting, population-genetic summaries, and conversion to downstream formats. These repeated scans are a recurring cost in post-calling analysis, and each is typically carried out by a tool that parses the entire record even when the operation depends on only one or two of its fields.

This inefficiency follows directly from the VCF design. A VCF record stores every property of a variant on a single line, beginning with the site-level fields (position, quality, filter status, and INFO annotations) and continuing, for every sample in the cohort, with a FORMAT block such as `GT:DP:GQ:AD` that encodes the genotype, read depth, genotype quality, and allelic depths. This layout makes VCF a general exchange format that any downstream program can read, which is why it has become the standard representation for variant data. It also means that a program answering a field-limited question, such as retaining sites with `QUAL` > 30 or measuring per-site missingness, must still decode the full record, including the per-sample values it will not use. On cohorts of thousands of samples, where the per-sample FORMAT block is by far the largest part of each record, decoding these unused fields typically dominates the running time of the operation.

Established tools take different routes to this problem. VCFtools (Danecek et al., 2011) parses the full text record. bcftools with HTSlib (Danecek et al., 2021; Bonfield et al., 2021) unpacks BCF fields in tiers and, given a tabix or CSI coordinate index (Li, 2011), restricts reading to requested genomic regions, so it avoids decoding data it does not need for site-level BCF queries. Fast parsing libraries such as cyvcf2 (Pedersen and Quinlan, 2017) likewise provide lazy access to individual fields, and dedicated engines re-encode genotypes into query-optimised stores for fast field- or sample-limited access, including GQT (Layer et al., 2016) and the fixed-schema binary genotypes of PLINK (Chang et al., 2015). Closest to the present work are expression-driven filters that evaluate user-written predicates directly against VCF and BCF records, including vembrane (Hartmann et al., 2023) and vcfexpress (Pedersen and Quinlan, 2025), the latter also implemented in Rust. These tools show that a compact expression language can express most filtering intent without bespoke code, and they establish the interface convention VariantFlow adopts.

What remains open is what the engine does with that expression once it has it. In the tools above the predicate selects records; it does not determine how much of each record is decoded, nor which parts of the file are read at all. VariantFlow derives both from the predicate: the set of fields decoded per record is the set the predicate references, and per-block summaries of those same fields allow whole compressed blocks to be skipped on non-positional criteria such as filter status or an allele-frequency threshold, which a coordinate index cannot evaluate.

VariantFlow addresses this inefficiency through selective execution. For each filter, statistic, or projection, it determines the minimal set of fields on which the result depends and decodes only those fields while streaming once through the file, leaving the remainder of each record unparsed; where the operation allows, it returns the original record text unchanged. VariantFlow applies this strategy to VCF/BCF filtering, VCFtools-compatible population-genetic summaries, and columnar export, and it reports a performance gain only where its output matches an established reference tool. It is intended to complement, not replace, bcftools, HTSlib, VCFtools, and GATK (McKenna et al., 2010).

## 2 Materials and Methods

### 2.1 Selective execution

Selective execution is the central design principle of VariantFlow. A VCF record stores every field for every sample on a single line, so a conventional parser must walk the whole line before it can return any part of it. Most analyses, however, depend on a small fraction of that line: a quality filter needs QUAL, a missingness summary needs only the genotype of each sample, and an allele-frequency calculation needs neither read depths nor genotype qualities. The remaining bytes are decoded and discarded.

Each query, whether a filter expression, a population-genetic statistic, or a column projection, is therefore compiled into a typed predicate tree that identifies the specific fields on which the result depends (Figure 1). VariantFlow then streams once through the file and, for each record, decodes only those fields. Rather than materialising the whole record as strings, it treats each record as a borrowed byte-slice view and locates the required site, INFO, and FORMAT fields by scanning the raw bytes in place, so no allocation is made for a field that is never read. Because the skipped fields cannot, by construction, affect the result, the output is identical to that of a tool that decodes every field. That equivalence is a design property rather than an approximation: VariantFlow reads fewer fields, never fewer records, and never samples or estimates. It is verified empirically in the Results.

**Figure 1.**
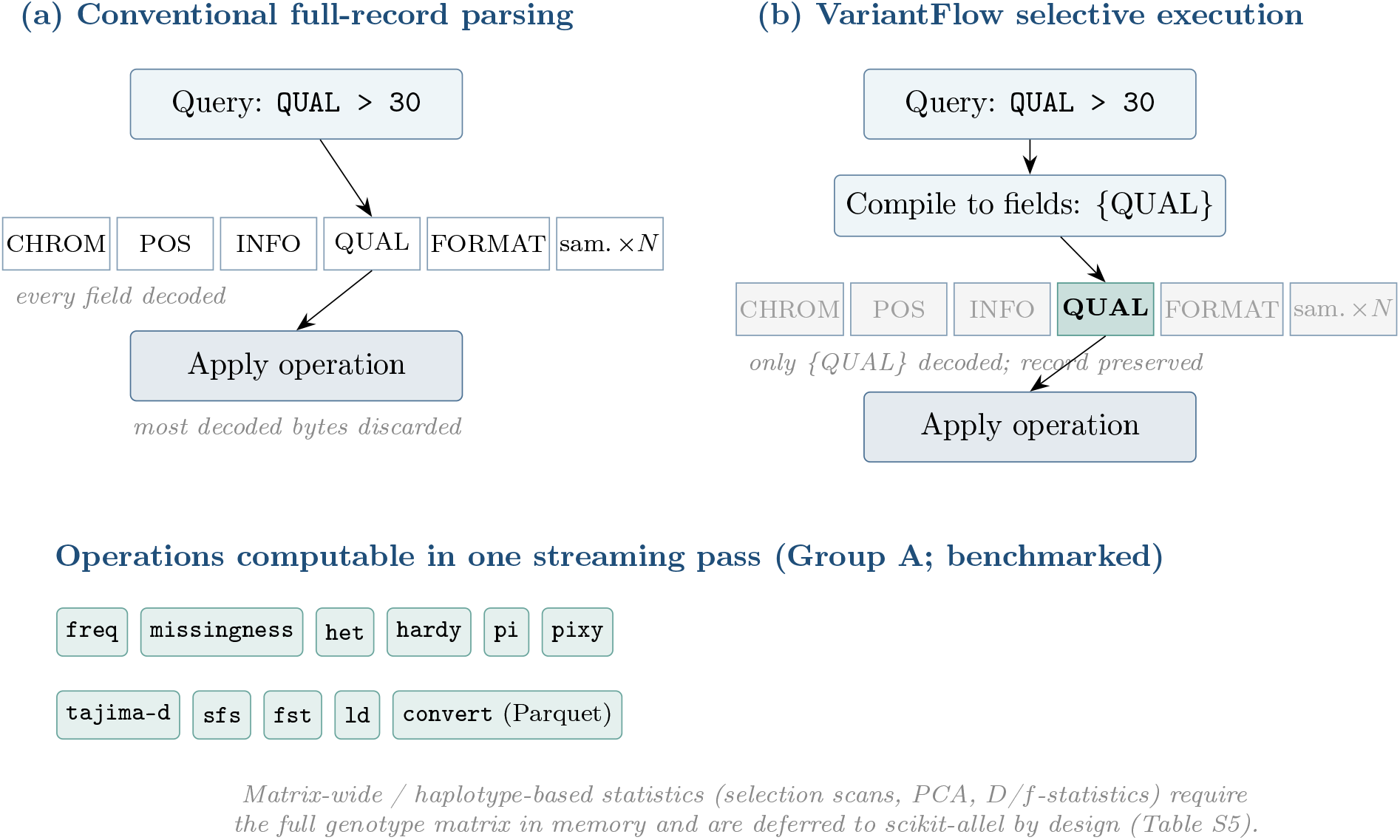
Selective execution versus conventional full-record parsing. (a) Conventional tools decode every field of every VCF record before applying an operation, so most decoded bytes are discarded. (b) VariantFlow compiles each query to the minimal set of fields it touches and decodes only those, preserving the original record. The lower band lists the population-genetics statistics and the streaming Parquet export that VariantFlow computes in a single pass; matrix-wide and haplotype-based statistics are deferred to scikit-allel (Supplementary Table S5).

### 2.2 Field-selective parsing of per-sample data

The cost that selective execution avoids grows with cohort size, because the per-sample block is the part of a VCF record that scales. In a 3 202-sample cohort each record carries thousands of `GT:DP:GQ:AD` entries, and a filter on read depth alone must otherwise traverse all of them. VariantFlow parses the FORMAT schema once per record, caches the position of the requested key such as `DP`, and reads that value by offset across all samples, avoiding a repeated scan of the layout for every sample. In its line-preserving mode, records that pass a filter are written in input order with their original text unchanged, so a filtered file remains byte-comparable with its source.

### 2.3 Compressed input, parallelism, and query-aware indexing

Compressed input is read through a native block-gzip (BGZF) reader with a worker-thread count capped automatically to the host. CPU-bound predicates are evaluated in ordered batches so that output order is preserved regardless of the number of threads.

For filters that are applied repeatedly and reject most records, an optional `.vfi` sidecar index stores per-block metadata: the genomic range, `QUAL` bounds, the set of `FILTER` values present, and summaries of selected INFO fields. The query planner consults this metadata and skips an entire compressed block whenever it proves that no record within it can satisfy the filter. The distinction from a coordinate index such as tabix (Li, 2011) is that the skipping criterion need not be positional. A tabix or CSI index answers “which blocks overlap this region”; it cannot answer “which blocks contain a record whose `FILTER` is `PASS`” or “whose allele frequency exceeds a threshold”, because those properties are not functions of genomic position. Storing per-block summaries of exactly the fields that predicates reference is what makes non-positional skipping possible, and it is the reason the largest speedups in this study arise for predicates that exclude most of the file. When few blocks can be excluded, the planner falls back to a sequential scan.

Through an optional HTSlib backend (Bonfield et al., 2021), VariantFlow additionally reads BCF (the binary encoding of VCF), restricts reading to a single indexed region, and writes BGZF output.

### 2.4 Population-genetic statistics and columnar export

For population-genetic analysis, the VCFtools-compatible subcommands operate on compact diploid biallelic genotype summaries and reproduce VCFtools output for allele frequency, missingness, heterozygosity, Hardy–Weinberg equilibrium, nucleotide diversity, Tajima’s *D*, linkage disequilibrium, and *F*_*ST*_ . Each is computed in a single streaming pass over sites or windows, so peak memory is set by the number of samples rather than the number of records. For repeated analytical queries, VariantFlow exports typed columns in the Parquet format (Apache Parquet Project, 2026), which columnar engines such as DuckDB (DuckDB Foundation, 2026) can query efficiently without re-parsing the VCF.

### 2.5 Interface and usage

The command-line interface follows the compositional, single-purpose style common in genomics, exposing the operations as a small tree of filtering, indexing, input/output, and population-genetic subcommands (Figure 2).

**Figure 2.**
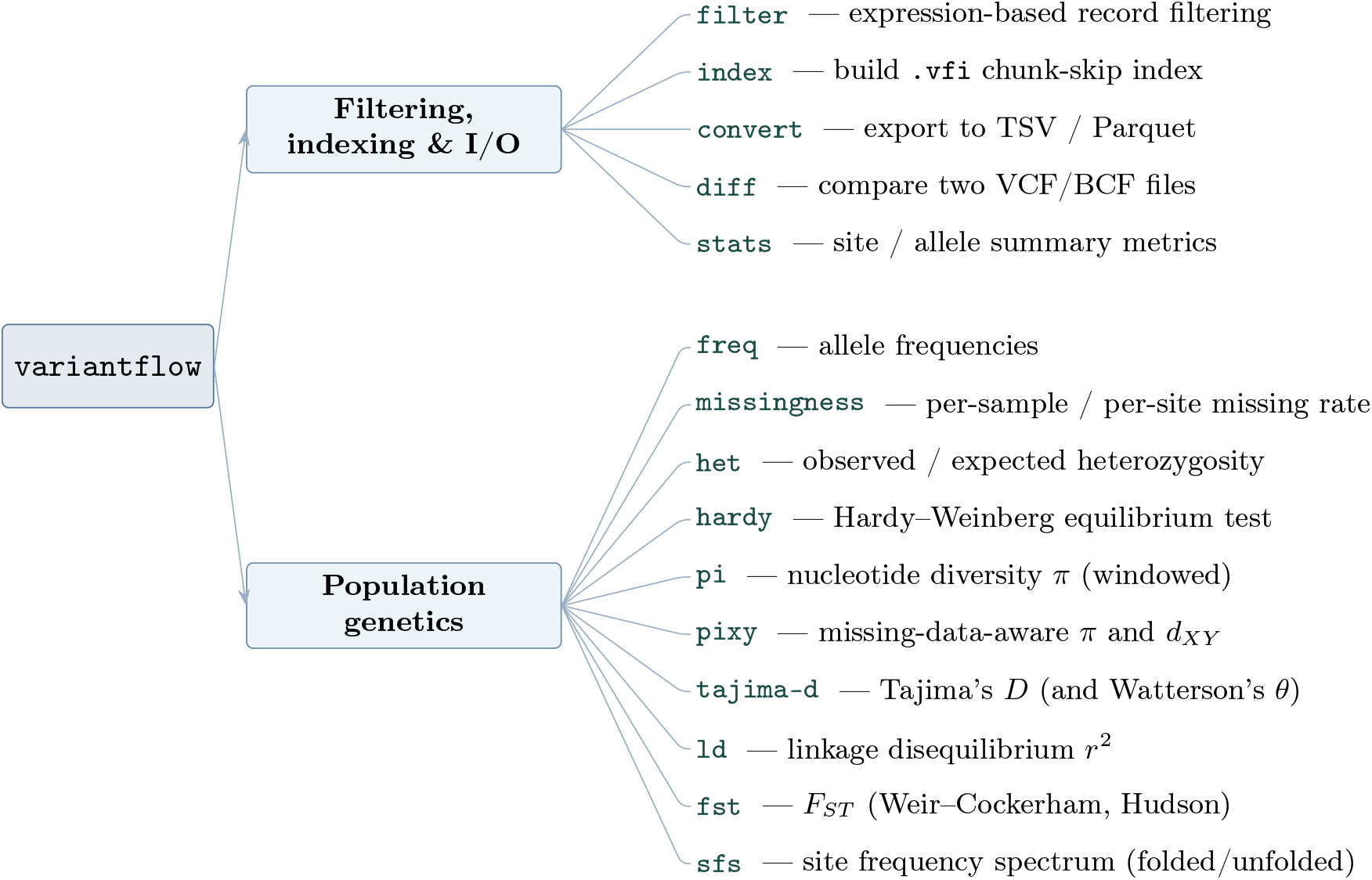
The VariantFlow command tree. Subcommands are grouped into two families that reflect the two ways selective execution is applied. The filtering, indexing, and input/output commands select records and fields, and return VCF, BCF, or Parquet; the population-genetics commands consume the same selective reader but return per-site or per-window statistics as tab-separated tables. The split matters in use because the first family composes with established tools in a pipeline, while the second terminates it. Each leaf gives the command and a one-line description of its function.

A representative FORMAT-aware filter keeps sites that pass a quality threshold and have at least one sample sequenced to sufficient depth:

~~~
variantflow filter chr22.vcf.gz \
–where “QUAL > 30 && ANY(FORMAT/DP > 20)” -o chr22.filtered.vcf
~~~

The output is a VCF whose passing records are byte-identical to the corresponding input lines. A two-population *F*_*ST*_ computation takes two files of sample identifiers and writes a tab-separated table:

~~~
variantflow fst chr22.vcf.gz –pop AFR.txt –pop EUR.txt -o fst.tsv
~~~

Each row of `fst.tsv` gives the chromosome, position, and the Weir–Cockerham estimate for that site, so a value of, for example, 0.31 at a given position indicates that roughly a third of the genetic variance there is partitioned between the two populations rather than within them. The same invocation pattern applies to the other summaries, which differ only in the statistic named and the options it requires. A worked end-to-end analysis of 1000 Genomes chromosome 22, from installation through interpretation, is provided as a supplementary user guide.

VariantFlow is written in Rust and builds from source with Cargo on Linux and macOS; prebuilt binaries and a container image are distributed with each release. The optional HTSlib backend is enabled at compile time and is not required for any result reported here.

### 2.6 Benchmark and validation protocol

All comparisons reported here are drawn from benchmark reports tracked in the repository, each recording the exact commands, the competitor version, and the correctness check applied. A speedup is reported only for an operation whose output has first been verified against a reference tool (Supplementary Table S1); operations that trail a competitor are treated as optimisation targets rather than claims, and are reported as measured.

Population-genetic benchmarks were run on an Apple M5 Max workstation and the index-assisted and columnar benchmarks on a Docker/Linux host. Within every comparison both tools were run on the same host, so no ratio spans machines. The full system configuration, tool versions, and dataset shapes are given in Supplementary Table S2. Timing is single-threaded wall-clock; memory is peak resident set size, and correctness was assessed by normalising both outputs and comparing them field by field.

## 3 Results

### 3.1 Selective filtering and query-aware indexing

On a 3 715-sample chromosome 22 call set aligned to the T2T-CHM13 reference (Nurk et al., 2022) and distributed by DDBJ, filters over FORMAT-rich predicates produced the same records as bcftools while running 4.74–17.78x faster (Table 1). These filtering measurements were taken on bounded subsets of 1 000, 10 000, and 50 000 records streamed from the 27 GB source file, which the harness does not cache locally; the population-genetic measurements reported below are whole-chromosome. Query-aware indexing with the `.vfi` sidecar gave the largest gains where most blocks can be excluded in advance. On the 1000 Genomes chromosome 22 data, a FILTER = PASS predicate, for which block metadata proves that no record in a skipped block can pass, ran 123–273x faster (0.213 s versus 58.03 s at the one-million-record tier), and a block-level allele-frequency threshold predicate ran 4.70–4.93x faster. The allele-frequency path trades memory for speed, reaching a 66.1 MB peak resident set at the one-million-record tier against bcftools’ 4.4 MB. When few blocks can be excluded, VariantFlow falls back to a sequential scan and the index confers no advantage; on synthetic datasets in which the planner cannot exclude blocks, the indexed path is slower than the unindexed one.

**Table 1.**
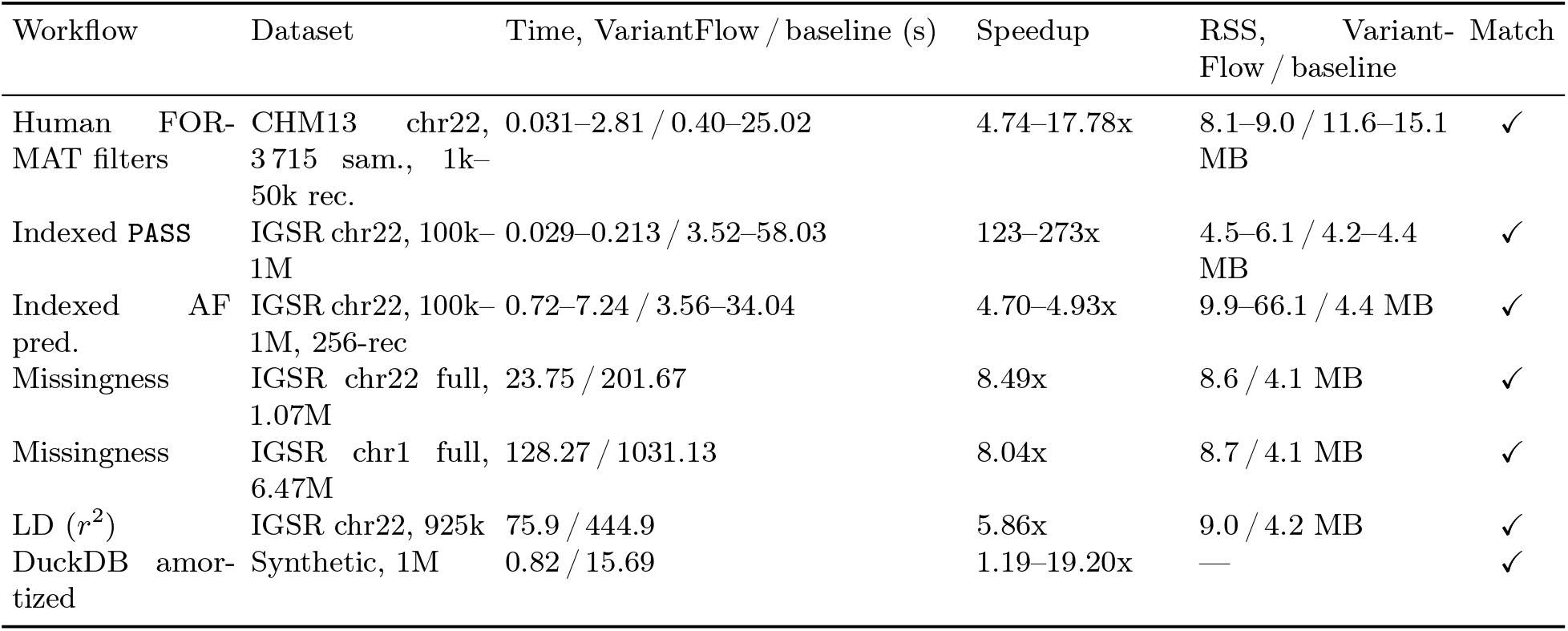
Correctness-matched benchmarks from tracked reports. Each row records exact commands, competitor version, correctness checks, and peak resident memory. IGSR rows use the 1000 Genomes 3202-sample high-coverage dataset. ✓ = output verified against the baseline: byte-identical for the population-genetic rows, and identical filtered core records for the filtering and indexed rows. Baselines: `bcftools filter` (filtering and indexed rows), VCFtools 0.1.17 (population summaries), repeated scans of the source file (DuckDB). Memory is peak resident set size; hosts differ by row and are given in Supplementary Table S2. See Supplementary Table S1 for cross-tool validation including scikit-allel.

### 3.2 Population-genetic summaries at whole-chromosome scale

Population-genetic summaries on the full 1000 Genomes 3 202-sample high-coverage cohort (Byrska-Bishop et al., 2022) confirm that selective execution scales to whole-chromosome data. Missingness computation, for which VariantFlow emits per-site and per-individual rates in a single pass, ran 8.49x faster than VCFtools on chromosome 22 (1,073,369 records; 23.75 s versus 201.67 s) and 8.04x faster on chromosome 1 (6,468,094 records; 128.27 s versus 1031.13 s), at a peak resident memory of 8.6 MB that was independent of chromosome size. The chromosome 1 summary therefore completes in about two minutes where VCFtools requires about seventeen. Bounded-distance linkage disequilibrium (*r*^2^, `–max-distance 500`) ran 5.9x faster on a 925,730-record dataset (75.9 s versus 444.9 s). The remaining VCFtools-compatible summaries were faster by smaller margins on the same cohort: allele frequency 3.68x, heterozygosity 1.46x, Tajima’s *D* 1.27x, Hardy–Weinberg equilibrium 1.23x, Weir–Cockerham *F*_*ST*_ 1.09x, and per-site *π* 1.08x. Selective execution therefore yields large gains where field selectivity is high and modest ones where nearly every genotype must be read regardless.

### 3.3 Columnar export for repeated queries

On a separate deterministic synthetic call set, exporting the data once to Parquet and issuing repeated queries through DuckDB was 1.19–19.20x faster than repeated scans of the source file, amortised over the query workload (Supplementary Table S3).

### 3.4 Correctness against established tools

Correctness was verified for every reported statistic, because the value of a faster tool depends on its returning the established answer. Allele frequency, per-site and per-individual missingness, nucleotide diversity (*π*), linkage disequilibrium, and Weir–Cockerham *F*_*ST*_ were byte-identical to VCFtools 0.1.17 on the same inputs (Supplementary Table S1); heterozygosity, Hardy–Weinberg equilibrium, and Tajima’s *D* are computed against the same VCFtools reference. An independent reimplementation in scikit-allel (Miles et al., 2024) reproduced per-individual missingness exactly across 1,066,557 variants and 3,202 samples (maximum absolute difference zero), and VariantFlow’s site-frequency spectrum matched scikit-allel’s folded and unfolded spectra bin for bin. Missing data is the regime in which diversity estimators are most likely to become biased, so it was tested directly. VariantFlow implements the unbiased per-site estimators of nucleotide diversity (*π*) and divergence (*d*_*XY*_) introduced by pixy (Korunes and Samuk, 2021), defined in Supplementary Note S1; on an all-sites cohort with 15% of genotypes set to missing, its per-window *π* and *d*_*XY*_ pairwise counts were identical to pixy across every window (Supplementary Table S4). The full validation procedure is described in Supplementary Note S2.

### 3.5 Scope of the implemented statistics

VariantFlow implements the population-genetic statistics that can be computed in a single streaming pass over sites or windows: allele frequency, missingness, heterozygosity, Hardy–Weinberg equilibrium, *π*, the missing-data-aware *π* and *d*_*XY*_ estimators, Tajima’s *D*, the site-frequency spectrum, *F*_*ST*_ , and linkage disequilibrium (Figure 1). Statistics that require the full genotype or haplotype matrix to be resident in memory, including selection scans, principal-component analysis, and *D*- and *f* -statistics, fall outside this streaming scope and are deferred to matrix-oriented libraries such as scikit-allel; Supplementary Table S5 maps the full correspondence.

## 4 Discussion

VariantFlow is a complement to the established tools rather than a replacement, and it is most effective for a defined class of tasks, namely selective filtering of VCF and BGZF files, index-assisted filtering that skips large regions of a file, the population-genetic summaries that VCFtools supports, and an export-once, query-many workflow based on Parquet. Its scope relative to existing tools is documented in a claim matrix distributed with the software, which records where it exceeds, matches, or complements each of them.

Different tools occupy complementary roles, and the appropriate choice depends on the task. bcftools remains the reference for general filtering, normalisation, and format conversion; VCFtools for established population-genetic file operations; DuckDB for ad hoc analytical queries over exported tables; and scikit-allel for matrix- and haplotype-based statistics such as selection scans, principal-component analysis, and *D*- and *f* -statistics, which require every sample’s genotypes to be held in memory simultaneously. PLINK (Chang et al., 2015) remains the reference for genome-wide association and matrix statistics computed on fixed-schema binary genotypes, and the cyvcf2 library (Pedersen and Quinlan, 2017) provides fast lazy VCF parsing for custom analyses in Python; VariantFlow differs in operating directly on VCF/BCF with query-driven field selection and byte- exact VCFtools-compatible output from a standalone command-line tool. VariantFlow occupies the gap between these, providing fast field-selective filtering and streamable per-site and per-window statistics on large cohorts, with export-once handoff to DuckDB and explicit deferral to scikit-allel for matrix-wide analyses (Supplementary Table S5). In practice, VariantFlow is the appropriate choice when interactive turnaround on large VCF files using a single workstation is the priority, and an established tool when its full generality is required.

GPU-accelerated pipelines such as NVIDIA Clara Parabricks target variant calling and the upstream processing that produces a VCF, using massive parallelism (NVIDIA Corporation, 2026). VariantFlow addresses a distinct bottleneck, the post-calling selection of fields on the commodity CPUs available in most laboratories, and the two approaches are complementary.

If selective execution becomes common in post-calling tools, routine analyses of large cohorts could move from batch to interactive timescales, and the export-once pattern could reduce redundant reading and rewriting of data. The groups most likely to benefit are those that repeatedly process large cohort VCFs on standard hardware, including population and conservation genomics, which compute diversity, differentiation, and linkage disequilibrium; plant and animal breeding programmes that screen large panels; large-cohort human-genomics quality control based on missingness and allele-frequency summaries; and developers who build scriptable filtering and export steps into pipelines. For these users the distinctive contribution of VariantFlow is that whole-cohort post-calling summaries become tractable on the hardware they already have. A whole-chromosome missingness summary that takes VCFtools 1031 s on chromosome 1 completes in 128 s on the same workstation, with byte-identical output.

VariantFlow has clear limits. It decodes only a defined subset of per-sample FORMAT fields, so analyses that depend on FORMAT keys it does not yet parse must use the established tools. Its index-assisted allele-frequency path trades memory for speed, reaching a 66 MB peak resident set at the one-million-record tier against bcftools’ 4.4 MB, and on datasets where few blocks can be excluded in advance the index confers no advantage over a sequential scan.

## 5 Conclusions

VariantFlow shows that deriving the decode set from the query itself, rather than parsing every field of every record, is sufficient to move whole-cohort post-calling analysis onto a single workstation while leaving results unchanged. The gains follow the mechanism rather than the implementation language. They are large where a query touches few fields or where block metadata excludes most of a file, and small where nearly every genotype must be read regardless. Reporting both regimes is what allows the principle to be judged rather than merely asserted.

For practitioners the immediate consequence is that summaries which previously ran as batch jobs can be run repeatedly while a question is still being formulated, without a compute cluster and without departing from the outputs established tools produce. Extending the expression language and the set of supported FORMAT fields is the principal direction of future work, as is broadening the per-block summaries that make non-positional skipping possible.

## Supporting information

User-Guide

## Author contributions

Following CRediT, E.E.: conceptualisation, software, validation, investigation, data curation, and writing of the original draft. J.-F.M.: supervision and writing through review and editing. Both authors read and approved the final manuscript.

## AI usage disclosure

AI coding assistants (OpenAI Codex and Anthropic Claude Code) were used for planning, code-review support, test scaffolding, benchmark-report organisation, and prose drafting support. The authors reviewed all output and remain responsible for the manuscript, code, benchmarks, and all scientific claims.

## Acknowledgements

Computing resources provided by SNIC/HPC2N at Umeå University.

## Data Accessibility and Benefit-Sharing Statement

### Data Accessibility Statement

VariantFlow’s source code, build instructions, and all reproducibility materials (exact commands, benchmark reports, and correctness-validation scripts) are openly available at https://github.com/ehsanestaji/VariantFlow under the MIT OR Apache- license. The version described in this manuscript (v1.5.0) is permanently archived at Zenodo, doi:10.5281/zenodo.21198172; the concept DOI doi:10.5281/zenodo.21198171 resolves to the latest release. Benchmark reports written after the v1.5.0 release, including the reproduction and cross-check reports cited in the supplementary material, are maintained on the repository’s `main` branch rather than in the frozen v1.5.0 archive. No new biological samples or sequence data were generated for this study. The benchmarks reuse publicly available, previously deposited variant call sets: the 1000 Genomes Project high-coverage release (International Genome Sample Resource) for the population-genetic and indexed-filter benchmarks, and the 3 715-sample chromosome 22 joint call set aligned to the T2T-CHM13 reference for the FORMAT-filter benchmark. The latter is a derived joint call over previously published public cohorts, distributed by the DNA Data Bank of Japan as files rather than as a deposited study, and therefore carries no study accession; it is identified by its stable distribution URL, https://ddbj.nig.ac.jp/public/public-human-genomes/CHM13/JointCall/CHM13_autosome_PAR.chr22.vcf.gz (server `Last-Modified` 2025-12-20T17:43:58Z; 27,232,829,080 bytes; CSI index alongside). All datasets are used under their existing public data-access terms. The Parquet and DuckDB benchmarks use a deterministic synthetic call set generated by a script in the repository. VariantFlow is written in portable Rust and builds from source with Cargo; platform-specific build notes and the optional HTSlib backend are described in the repository README.

### Benefit-Sharing Statement

This study did not involve the collection of new biological specimens, field sampling, or engagement with Indigenous or local communities; it reuses previously published, de-identified public human genomic datasets. Benefit-sharing provisions are therefore not applicable to this work.

## Supplementary material

The supplementary document (`supplementary.pdf`) contains the following items; each is self-contained and begins on its own page:

- Supplementary Table S1 — cross-tool correctness validation against VCFtools and scikit-allel.
- Supplementary Table S2 — benchmark system configuration.
- Supplementary Table S3 — timing and memory for the streaming population-genetics and Parquet/DuckDB workflows.
- Supplementary Table S4 — missing-data validation against pixy.
- Supplementary Table S5 — statistics coverage relative to scikit-allel, grouped by computational class.
- Supplementary Note S1 — missing-data-aware *π* and *d*_*XY*_ .
- Supplementary Note S2 — correctness validation methodology.

An end-to-end population-genetics tutorial (1000 Genomes chromosome 22, from installation to interpretation) is provided in the software repository.

