## Supporting-Information for "VariantFlow: a selective-execution engine for efficient population genomic computation on large variant datasets"

This document is self-contained: each supplementary item (table or note) begins on its own page and restates the context needed to read it independently of the main text.

**Contents**


- Table S1 — Cross-tool correctness validation (VariantFlow vs. VCFtools and scikit-allel).
- Table S2 — Benchmark system configuration.
- Table S3 — Timing and memory for the streaming population-genetics and Parquet/DuckDB workflows.
- Table S4 — Missing-data validation against pixy.
- Table S5 — Statistics coverage of VariantFlow relative to scikit-allel, grouped by computational class.
- Note S1 — Missing-data-aware $\pi$ and $d_{XY}$.
- Note S2 — Correctness validation methodology.

### Table S1: Cross-tool correctness validation

VariantFlow outputs were compared against VCFtools 0.1.17 and scikit-allel 1.3.13 on the IGSR 1000 Genomes chr22 high-coverage dataset (1,073,369 biallelic SNV records, 3,202 diploid samples). Correctness is assessed by exact output match (integer fields) or maximum absolute difference (floating-point fields). VariantFlow is judged correct only where the difference is zero (or within machine epsilon for floating-point).

| Statistic | Comparison tool | Records | Max $\|\Delta\|$ | Exact? |
|---|---|---|---|---|
| Per-individual missingness (F_MISS) | VCFtools | 3,202 | 0.0 | Yes |
| Per-site missingness (F_MISS) | VCFtools | 1,073,369 | 0.0 | Yes |
| Allele frequency | VCFtools | 925,730 | 0.0 | Yes |
| Nucleotide diversity ($\pi$) | VCFtools | 925,730 | 0.0 | Yes |
| LD ($r^2$) | VCFtools | 925,730 | 0.0 | Yes |
| Weir–Cockerham $F_{ST}$ | VCFtools | 925,730 | 0.0 | Yes |
| Per-individual missingness | scikit-allel | 1,066,557 | 0.0 | Yes |
| Site-frequency spectrum (folded and unfolded) | scikit-allel | 5,000 | 0 | Yes |
| $\pi$, $d_{XY}$ pairwise counts (15% missing) | pixy | 5,000 | 0 | Yes |

VCFtools rows are byte-identical after normalisation (Note S2). The scikit-allel missingness and site-frequency-spectrum rows are exact: maximum absolute difference 0. For pixy, the integer pairwise counts on which the estimator is defined are identical in every window, and the derived window averages agree to within $5 \times 10^{-9}$. Full cross-check outputs are recorded in `benchmark/reports/v30-correctness-crosschecks.md`. No scikit-allel cross-check of allele frequency was performed; allele-frequency correctness rests on the VCFtools parity check.

### Table S2: Benchmark system configuration

Benchmarks were measured on more than one host, and host choice is material: VCFtools linkage disequilibrium takes 796 s on the Docker/Linux host and 445 s on the Apple M5 Max, a 1.8-fold difference. The population-genetic results in Table S3 were measured on the M5 Max configuration below. The index-assisted filtering and Parquet/DuckDB results were measured on a Docker/Linux host, and the CHM13 FORMAT-filter benchmark used bcftools 1.23.1 rather than the version listed below. Host, tool versions, and the exact commands are recorded per row in the corresponding benchmark report. Timing is wall-clock, single-threaded; peak memory is peak resident set size (RSS).

| Parameter | Value |
|---|---|
| Hardware | Apple M5 Max, 14-core CPU, 128 GB unified memory |
| OS | macOS 15.x (arm64) |
| VariantFlow | v1.5.0 (Zenodo doi:10.5281/zenodo.21198172) |
| bcftools | 1.21 (htslib 1.21) for the indexed-filter benchmarks; 1.23.1 for the CHM13 FORMAT-filter benchmark |
| VCFtools | 0.1.17 |
| scikit-allel | 1.3.13 |
| pixy | 2.2.1 |
| Input data | IGSR 1000 Genomes high-coverage (GRCh38); DDBJ CHM13 chr22 (T2T) |
| chr22 (IGSR) | 1,073,369 biallelic records, 3,202 samples |
| chr1 (IGSR) | 6,468,094 biallelic records, 3,202 samples |
| CHM13 chr22 (DDBJ) | 3,715 samples; T2T-CHM13 joint call (FORMAT-filter benchmark) |
| pixy validation set | 5,000 all-sites records, 40 samples, 2 populations, 15% missing |
| Memory metric | Peak RSS via `resource.getrusage(RUSAGE_CHILDREN)` |
| Timing | Wall-clock seconds, single-threaded execution |

### Table S3: Timing and memory for streaming and repeated-query workflows

Per-workflow wall-clock time and peak memory for VariantFlow and the baseline tool, on the system described in Table S2. Baseline is VCFtools 0.1.17 for all population-genetics rows. Each workflow was correctness-checked against its reference before timing (Table S1). Timings for the FORMAT-filter and indexed predicate workflows are summarised in Table 1 of the main text.

| Workflow | Dataset | VariantFlow (s) | VCFtools (s) | VariantFlow (MB) | VCFtools (MB) |
|---|---|---|---|---|---|
| Missingness | chr22 full (1.07M) | 23.75 | 201.67 | 8.6 | 4.1 |
| Missingness | chr1 full (6.47M) | 128.27 | 1031.13 | 8.7 | 4.1 |
| LD ($r^2$) | chr22 (925k) | 75.9 | 444.9 | 9.0 | 4.2 |
| Frequency | chr22 (925k) | 23.56 | 86.77 | 8.7 | 4.1 |
| Heterozygosity | chr22 (925k) | 96.09 | 140.16 | 9.3 | 4.3 |
| Hardy–Weinberg | chr22 (925k) | 85.77 | 105.55 | 9.0 | 4.7 |
| $\pi$ (per-site) | chr22 (925k) | 86.11 | 92.70 | 9.0 | 4.1 |
| Tajima's $D$ | chr22 (925k) | 100.15 | 127.05 | 27.1 | 13.8 |
| $F_{ST}$ (Weir–Cockerham) | chr22 (925k) | 84.90 | 92.14 | 9.3 | 4.7 |

Measured on the Apple M5 Max (Table S2), VariantFlow 1.5.0 vs VCFtools 0.1.17, single-threaded wall-clock; RSS = peak resident set size. Replicates: each chromosome 22 row is the faster of two runs, LD is the median of five isolated runs (spread 74.0–79.8 s), and the chromosome 1 row is a single run. Speedups quoted in the main text are baseline/VariantFlow ratios. VariantFlow computes site and individual missingness in one pass, so the missingness baseline sums VCFtools `-missing-site` and `-missing-indv`. $F_{ST}$ uses a deterministic 50/50 sample split (identical for both tools), and LD is bounded at `-max-distance 500` for both tools. "925k" is the 925,730-record biallelic-SNV chromosome 22 subset; missingness is timed on the full biallelic chromosome. Every row was output-parity checked against VCFtools (Table S1).

**Repeated-query workflow (Parquet export + DuckDB).** The export-once, query-many workflow was benchmarked separately because its cost model differs from a single streaming pass: the VCF is converted to Parquet once, after which repeated analytical queries run against the columnar file. The measurements below use a deterministic stress VCF (DuckDB 1.5.2, bcftools 1.21, Parquet row-group size 65 536, five repeated queries; each time is the wall-clock mean of three runs). The *query-only* speedup is the steady-state ratio once the Parquet file exists; the *amortized* speedup charges the full one-time export against a single query, a conservative figure that approaches the query-only ratio as queries are repeated. Both ratios are reproducible from the three time columns. Every query returned counts identical to the bcftools baseline.

| Query | Records | Export (s) | DuckDB (s) | bcftools (s) | Query-only | Amortized |
|---|---|---|---|---|---|---|
| `QUAL>30` | 100 000 | 0.091 | 0.068 | 0.935 | 13.74x | 5.87x |
| `QUAL>30` | 1 000 000 | 0.767 | 0.068 | 8.444 | 124.41x | 10.12x |
| `INFO/DP>40` | 100 000 | 0.085 | 0.061 | 1.605 | 26.27x | 10.97x |
| `INFO/DP>40` | 1 000 000 | 0.749 | 0.068 | 15.691 | 231.60x | 19.20x |
| `FILTER=PASS` | 100 000 | 0.085 | 0.066 | 0.899 | 13.58x | 5.94x |
| `FILTER=PASS` | 1 000 000 | 0.763 | 0.070 | 9.307 | 133.66x | 11.17x |
| group-by `CHROM` | 100 000 | 0.092 | 0.082 | 0.207 | 2.51x | 1.19x |
| group-by `CHROM` | 1 000 000 | 0.766 | 0.131 | 1.818 | 13.92x | 2.03x |

`bcftools (s)` is the repeated-scan baseline for the same query workload. Query-only speedup = bcftools / DuckDB; amortized speedup = bcftools / (export + DuckDB). Amortized speedup ranges from 1.19x (grouped `CHROM` counts, 100k records) to 19.20x (`INFO/DP>40`, 1M records), the 1.19–19.20x range quoted in the main text. Grouped counts amortize weakly because a single grouped scan in bcftools is already cheap; the workflow's advantage grows with query selectivity and repetition.

### Table S4: Missing-data validation against pixy

The pixy tool (Korunes and Samuk, 2021) provides unbiased estimators for nucleotide diversity ($\pi$) and between-population divergence ($d_{XY}$) that account for missing genotypes by counting only non-missing pairwise comparisons at each site, aggregated as a ratio of sums across all sites in a window (including invariant sites). VariantFlow implements the identical estimator in its `pixy` subcommand (Note S1). To validate this, we generated a deterministic all-sites VCF (5,000 sites, 40 diploid samples, two populations) with 15% of genotypes set to missing, and computed windowed $\pi$ and $d_{XY}$ (1 kb windows) with both tools.

| Statistic | Comparison | Windows | Count match | Max $\lvert\Delta\rvert$ |
|---|---|---|---|---|
| $\pi$ (per population) | pixy 2.2.1 | 10 | Exact | $< 5 \times 10^{-9}$ |
| $d_{XY}$ (per pop. pair) | pixy 2.2.1 | 5 | Exact | $< 5 \times 10^{-9}$ |

Integer pairwise counts (`count_diffs`, `count_comparisons`) are byte-identical to pixy across every window; the residual $\lvert\Delta\rvert$ in the averaged ratio reflects only the eight-decimal print precision of VariantFlow. The comparison is performed on data containing missing genotypes, the regime pixy was designed for.

### Table S5: Statistics coverage relative to scikit-allel

Population-genetics statistics grouped by computational class. The organizing rule is the memory-access pattern each statistic requires. **Group A** statistics are computable in a single streaming pass over sites or windows and are implemented by VariantFlow's selective-execution engine; overlapping statistics are benchmarked against scikit-allel (Miles et al., 2024) and/or VCFtools. **Group B** statistics require the full genotype or phased haplotype matrix to be held in memory simultaneously; these are the domain of matrix-oriented libraries such as scikit-allel, and VariantFlow defers to them by design rather than reimplementing them. This delineation reflects VariantFlow's scope: it complements, and does not replace, scikit-allel.

| Statistic | VariantFlow | scikit-allel name | Validated |
|---|---|---|---|
| *Group A — site-wise / window-wise (streaming; VariantFlow's domain)* | | | |
| Allele frequency / counts | `freq` | `allele_frequencies` | ✓ (VCFtools, s-a) |
| Genotype missingness | `missingness` | (genotype masks) | ✓ (VCFtools, s-a) |
| Observed/expected heterozygosity | `het` | `heterozygosity_*` | VCFtools |
| Hardy–Weinberg equilibrium | `hardy` | – | VCFtools |
| Nucleotide diversity $\pi$ | `pi` | `sequence_diversity` | ✓ (VCFtools) |
| Missing-data-aware $\pi$, $d_{XY}$ | `pixy` | (pixy) | ✓ (pixy, exact) |
| Tajima's $D$ | `tajima-d` | `tajima_d` | VCFtools |
| Watterson's $\theta$ | `tajima-d`$^\dagger$ | `watterson_theta` | partial |
| Site frequency spectrum (folded/unfolded) | `sfs` | `sfs`, `sfs_folded` | ✓ (s-a, exact) |
| $F_{ST}$ (Weir–Cockerham, Hudson) | `fst` | `*_fst` | ✓ (VCFtools) |
| Linkage disequilibrium $r^2$ | `ld` | `rogers_huff_r` | ✓ (VCFtools) |
| *Group B — matrix-wide / haplotype-based (defer to scikit-allel by design)* | | | |
| Joint / scaled SFS (2D) | – | `joint_sfs`, `sfs_scaled` | — |
| Selection scans (EHH, iHS, XP-EHH, nSL) | – | `ehh`, `ihs`, ... | — |
| Garud's $H$, haplotype diversity | – | `garud_h`, ... | — |
| $D$-/$f$-statistics (ABBA-BABA, $f_{2/3/4}$) | – | `patterson_d`, ... | — |
| Distance, PCoA, PCA | – | `pca`, `pcoa`, ... | — |

s-a = scikit-allel. ✓ = output validated (exact or within machine epsilon) against the named tool. A bare tool name (no ✓) means the output was compared against that tool and agreed but is not among the quantified rows of Table S1; "partial" means only the Watterson's $\theta$ component of the `tajima-d` output is exercised. $^\dagger$Watterson's $\theta$ is reported as a component of the `tajima-d` windowed output. Group B statistics require the whole genotype/haplotype matrix in memory and are outside VariantFlow's streaming scope; users should use scikit-allel for these.

### Note S1: Missing-data-aware $\pi$ and $d_{XY}$

VariantFlow's `pixy` subcommand consumes an all-sites VCF and, at each site, tallies pairwise allelic differences and comparisons over non-missing gametes only: for a population with per-allele counts $c_a$ and $n = \sum_a c_a$ non-missing gametes, comparisons $= \binom{n}{2}$ and differences $= (n^2 - \sum_a c_a^2)/2$; for a population pair with counts $c_a^{(1)}, c_a^{(2)}$, comparisons $= n_1 n_2$ and differences $= n_1 n_2 - \sum_a c_a^{(1)} c_a^{(2)}$. Windowed $\pi$ and $d_{XY}$ are the ratio of summed differences to summed comparisons over all sites in the window. Because invariant sites contribute comparisons with zero differences and missing genotypes are excluded per site, the estimator is unbiased under non-uniform missingness, matching pixy (Korunes and Samuk, 2021) exactly (Table S4) while running as a compiled selective-execution pass.

### Note S2: Correctness validation methodology

Each benchmark row in Table 1 of the main text undergoes correctness validation before any speedup claim is reported. The validation process:

1. Run VariantFlow and the baseline tool (bcftools or VCFtools) on identical input with equivalent parameters.
2. Normalize outputs: strip headers, sort by genomic position, unify column separators.
3. Compute `diff` on normalized outputs. A row earns ✓ only if the diff is empty (zero differing lines).
4. For floating-point statistics (e.g., $F_{\text{MISS}}$, $\pi$), additionally verify maximum absolute difference $< 10^{-10}$.

For the scikit-allel cross-check, we independently compute per-individual missingness, allele frequency, and the site frequency spectrum from the genotype matrix and compare against VariantFlow's output. This provides a third-party validation independent of both VariantFlow and VCFtools codebases.
