## Supplementary material for "VariantFlow: a selective-execution engine for efficient population genomic computation on large variant datasets": User-Guide

### Supplementary User Guide: an end-to-end population-genetics analysis of 1000 Genomes chromosome 22 with VariantFlow


**Abstract**

This supplementary note is a beginner-friendly, end-to-end walkthrough of a population-genetics analysis of human chromosome 22 from the 1000 Genomes Project using VariantFlow, a command-line tool for post-calling VCF operations. It is written for a reader with basic command-line familiarity but no prior bioinformatics experience. Every command is copy-pasteable; each step explains what it does and why. Slow steps include a small-region shortcut so the analysis can be reproduced quickly.


#### 1 Introduction

A **VCF** (Variant Call Format) file is the standard text table of genetic variants: each row is a genomic position where individuals differ, and each column after the fixed fields is one sample's genotype. VariantFlow performs *post-calling* VCF operations—the analyses that happen after variants have been called. Its guiding idea is **selective execution**: it reads and computes only the parts of the file a query needs, which keeps filtering and per-site statistics fast on large files. It reproduces the numbers of established tools (VCFtools, pixy, scikit-allel) from a single self-contained binary.

By the end of this guide you will have downloaded and indexed a real 1000 Genomes chromosome-22 VCF, reduced it to clean biallelic SNPs, and computed allele frequencies, missingness, heterozygosity, Hardy–Weinberg tests, nucleotide diversity ($\pi$), Tajima's D, linkage disequilibrium, and Fst between two human populations—and read what those numbers mean, cautiously. All commands are run as `variantflow <command>`.

**Time and disk warning.** The full chromosome-22 file is several gigabytes and some computations take a while. Wherever a step is slow, this guide offers a small-region shortcut (a $\sim$1 Mb slice) so you can follow along in minutes. Do the small-region version first; scale up once comfortable.

#### 2 Installation

##### 2.1 Build VariantFlow from source

VariantFlow is written in Rust. You need the Rust toolchain (`cargo`); if you do not have it:

```
# Install the Rust toolchain (rustup); accept defaults.
curl --proto '=https' --tlsv1.2 -sSf https://sh.rustup.rs | sh
source "$HOME/.cargo/env"
rustc --version
cargo --version
```

Build the release binary from the repository root:

```
# Standard build (pure-Rust VCF/BCF text path)
cargo build --release
./target/release/variantflow --version
```

For BCF input or region-based queries backed by HTSlib (e.g. `-region chr22:1-1000000` on a `.bcf`), build with the HTSlib feature:

```
cargo build --release --features htslib-static
```

Confirm the tool and list the subcommands:

```
./target/release/variantflow --help
```

You should see: `filter`, `stats`, `freq`, `missingness`, `hardy`, `het`, `fst`, `pi`, `pixy`, `tajima-d`, `ld`, `index`, `diff`, `convert`. For readability, the rest of this guide writes `variantflow`; either add the binary to `PATH` (`export PATH="$PWD/target/release:$PATH"`) or type the full path.

##### 2.2 Companion tools and dependencies

A few standard tools do the jobs VariantFlow deliberately does not (downloading, indexing, and heavy VCF surgery). The easiest route is Bioconda:

```
# One-time channel setup (order matters)
conda config --add channels bioconda
conda config --add channels conda-forge

# Companions in a fresh environment
conda create -n popgen -c bioconda -c conda-forge \
  bcftools htslib vcftools wget
conda activate popgen
```

Here `bcftools`/`htslib` provide `tabix` and `bgzip` (indexing, subsetting, slicing), `vcftools` enables optional cross-checks, and `wget` downloads data. Version-check everything:

```
bcftools --version
tabix --version
bgzip  --version
vcftools --version
wget --version | head -n1
```

#### 3 Getting the data

We use the 1000 Genomes Project 30× high-coverage phased panel (3202 samples) aligned to GRCh38, hosted by IGSR/EBI on EBI's public FTP. This file is **large** (several GB); Section 3.2 shows a 1 Mb slice for quick work.

##### 3.1 Download the full chromosome-22 panel

```
mkdir -p data && cd data

```

```
BASE="https://ftp.1000genomes.ebi.ac.uk/vol1/ftp/data_collections/1000
    G_2504_high_coverage/working/20220422_3202_phased_SNV_INDEL_SV"
FILE="1kGP_high_coverage_Illumina.chr22.filtered.SNV_INDEL_SV_phased_panel.vcf.gz"

# VCF and its tabix index (.tbi); -c resumes partial downloads
wget -c "${BASE}/${FILE}"
wget -c "${BASE}/${FILE}.tbi"
```

If the `.tbi` is absent, build it and check the file opens:

```
tabix -p vcf "${FILE}"
variantflow stats "${FILE}"
```

##### 3.2 Small-region shortcut (recommended first)

Slicing a 1 Mb window with `tabix` yields a small file that runs every downstream step in seconds (needs the `.tbi` index):

```
# Extract chr22:20,000,000-21,000,000
tabix -h "${FILE}" chr22:20000000-21000000 | bgzip > chr22.slice.vcf.gz
tabix -p vcf chr22.slice.vcf.gz
variantflow stats chr22.slice.vcf.gz
```

**Chromosome naming.** These files use `chr22` (with the `chr` prefix). If a region query returns nothing, check whether your file uses `22` instead and adjust (e.g. `22:20000000-21000000`). Set one variable so you can switch between the slice and the full file:

```
# Follow along fast with the slice:
VCF=chr22.slice.vcf.gz
# ...or run the full analysis:
# VCF="1kGP_high_coverage_Illumina.chr22.filtered.SNV_INDEL_SV_phased_panel.vcf.gz"
```

#### 4 Preprocessing

Raw call sets contain multiallelic sites, indels, and structural variants; most classic statistics assume **biallelic SNPs**.

##### 4.1 Reduce to biallelic SNPs (bcftools)

```
bcftools view -m2 -M2 -v snps "$VCF" -Oz -o chr22.snps.vcf.gz
tabix -p vcf chr22.snps.vcf.gz
```

Here `-m2 -M2` keeps sites with exactly two alleles (biallelic), `-v snps` keeps SNPs only, `-Oz` writes bgzip output, and `tabix` indexes it.

##### 4.2 Optional: select a subset of samples

Per-sample commands accept a plain-text file, **one sample ID per line** (blank lines and `#` comments ignored), via `-keep` (use only these) or `-remove` (drop these):

```
cat > keep.txt <<'EOF'
# one sample ID per line
HG00096
HG00097
HG00099
EOF
```

##### 4.3 Quality filtering with `filter`

`variantflow filter` selects records with a predicate passed to `-where`. The expression language understands `CHROM`, `POS`, `QUAL`, `FILTER`, `INFO/<TAG>` (with array indexing like `INFO/AF[1]`), and `FORMAT/<TAG>`, plus the aggregate `N_PASS(...)`.

```
# Keep only PASS sites with a decent quality score
variantflow filter chr22.snps.vcf.gz \
  --where 'QUAL > 30 && FILTER == "PASS"' \
  -o chr22.clean.vcf.gz
tabix -p vcf chr22.clean.vcf.gz

# Common variants only (alt-allele frequency above 5%)
variantflow filter chr22.snps.vcf.gz \
  --where 'INFO/AF[0] > 0.05' -o chr22.common.vcf.gz

# Restrict to a region while filtering (needs an index)
variantflow filter chr22.snps.vcf.gz \
  --region chr22:20000000-21000000 --where 'QUAL > 30' -o chr22.region.vcf.gz

# Require >=2 samples with a well-covered alternate allele
variantflow filter chr22.snps.vcf.gz \
  --where 'N_PASS(FORMAT/AD[1] > 10) >= 2' -o chr22.cohort.vcf.gz
```

**Note on this dataset.** The 1000 Genomes high-coverage panel is already a filtered, phased release, so most sites are `PASS` and `QUAL` is not always informative. The filter step matters more on your own freshly called data; here it mainly demonstrates the syntax. The important cleanup is the biallelic-SNP reduction. From here on, set:

```
SNPS=chr22.snps.vcf.gz
```

#### 5 Population-genetics calculations

Each subsection gives *what it measures*, *the command*, and *how to read the output*. Outputs are tab-separated tables.

##### 5.1 Allele frequency (`freq`)

**What it measures.** How common each allele is at every site—the building block for almost everything else.

```
variantflow freq "$SNPS" -o chr22.frq
```

Columns: `CHROM POS N_ALLELES N_CHR {ALLELE:FREQ}`. `N_CHR` is the number of chromosomes with data (twice the number of genotyped diploid samples). Restrict to a group with `-keep` / `-remove`.

##### 5.2 Missingness (`missingness`)

**What it measures.** How much genotype data is absent, per site and per individual. High missingness biases every other statistic.

```
variantflow missingness "$SNPS" -o chr22.miss
```

This writes `chr22.miss.lmiss` (per-site) and `chr22.miss.imiss` (per-individual). `F_MISS` is the fraction missing (0 = complete). On the phased panel it is essentially 0—a useful sanity check.

##### 5.3 Individual heterozygosity (`het`)

**What it measures.** Per-sample observed vs. expected homozygosity, summarised as an inbreeding coefficient $F$.

```
variantflow het "$SNPS" -o chr22.het
```

Columns: `INDV O_HOM E_HOM N_SITES F`. $F \approx 0$ matches Hardy–Weinberg expectation; $F > 0$ means fewer heterozygotes than expected (possible inbreeding or pooled structure); $F < 0$ means more (possible contamination or mixing groups).

##### 5.4 Hardy–Weinberg equilibrium (`hardy`)

**What it measures.** Whether observed genotype counts match Hardy–Weinberg proportions, with a chi-square test.

```
variantflow hardy "$SNPS" -o chr22.hwe
```

Columns include `OBS_HOM_REF OBS_HET OBS_HOM_ALT` and `CHISQ_HWE`. In a pooled multi-population sample, apparent departures are often *population structure* (the Wahlund effect), not genotyping error.

##### 5.5 Nucleotide diversity $\pi$ (`pi`, windowed)

**What it measures.** Average nucleotide differences per site between two random sequences—a core measure of within-population diversity, reported in windows.

```
variantflow pi "$SNPS" --window-size 100000 -o chr22.windowed.pi
```

Higher $\pi$ means a more diverse region. Use `-keep` to compute $\pi$ within a single population; $\pi$ mixed across populations is inflated by between-group differences.

##### 5.6 Tajima's D (`tajima-d`)

**What it measures.** Compares two diversity estimates to detect departures from neutral, constant-size evolution, per window.

```
variantflow tajima-d "$SNPS" --window-size 100000 -o chr22.tajimaD.tsv
```

$D \approx 0$ is consistent with neutrality; $D < 0$ (excess rare variants) with expansion or selection; $D > 0$ (excess intermediate-frequency variants) with balancing selection or a bottleneck. A value alone never proves selection.

##### 5.7 Linkage disequilibrium (ld)

**What it measures.** Non-random association between alleles at nearby sites, as a function of distance.

```
variantflow ld "$SNPS" --max-distance 50000 -o chr22.geno.ld
```

LD generally decays with distance. Keep `-max-distance` modest: the number of site pairs (and runtime) grows quickly.

##### 5.8 Fst between two populations (fst)

**What it measures.** Genetic differentiation between two populations: 0 = identical allele frequencies, higher = more differentiated.

First build two population files from the 1000 Genomes sample panel, which maps each sample to its population:

```
PED="https://ftp.1000genomes.ebi.ac.uk/vol1/ftp/data_collections/1000
    G_2504_high_coverage/20130606_g1k_3202_samples_ped_population.txt"
wget -c "$PED" -O samples.panel
head -n1 samples.panel   # inspect columns first!

# Extract two population lists (one sample ID per line).
# Adjust the awk field number to match the sample-ID column.
awk 'NR>1 && $0 ~ /GBR/ {print $2}' samples.panel > GBR.txt   # British
awk 'NR>1 && $0 ~ /YRI/ {print $2}' samples.panel > YRI.txt   # Yoruba
wc -l GBR.txt YRI.txt
```

Different panel files order columns differently; inspect the header and set the `awk` fields so each output is a plain list of sample IDs, one per line. Then compute Fst (Hudson estimator by default; `-estimator weir-cockerham` is also available):

```
variantflow fst "$SNPS" --pop GBR.txt --pop YRI.txt -o chr22.GBR_YRI.fst
```

Output columns: `CHROM POS HUDSON_FST` (per site). Fst $\approx 0$ means near-identical frequencies; human continental comparisons are often $\sim$0.1; high-Fst sites are selection candidates, not conclusions. Sites monomorphic in both groups return `nan` (undefined)—expected, not an error.

##### 5.9 Missing-data-aware $\pi$ and dxy (pixy)

**What it measures.** Windowed within-population diversity ($\pi$) and between-population divergence (dxy) using the pixy estimator, which stays unbiased under missing data by counting only non-missing pairwise comparisons.

**Key requirement:** `pixy` needs an **all-sites VCF**—one that includes invariant (monomorphic) sites, not just variant sites—because correct $\pi$/dxy is the ratio of differences to comparisons over all callable sites. The standard 1000 Genomes panel is variant-only, so this command targets an all-sites call set (e.g. produced with `bcftools mpileup`/`call` emitting all sites).

```
# populations file: two whitespace/tab separated columns per line:
#   SampleID    PopulationName
#   HG00096   GBR
#   NA18486   YRI
variantflow pixy allsites.vcf.gz \
  --populations pops.txt \
  --window-size 10000 \
  --out-pi   chr22.pixy_pi.tsv \
  --out-dxy  chr22.pixy_dxy.tsv
```

Note that `pixy` takes the input as a **positional** argument (not `-input`), and writes outputs to `-out-pi` and `-out-dxy` (there is no `-o`). Read `chr22.pixy_pi.tsv` (per-population $\pi$ per window) and `chr22.pixy_dxy.tsv` (per-pair dxy per window) as in the $\pi$ and Fst sections, with missing data handled correctly.

##### 5.10 Site-frequency spectrum (`sfs`)

**Verification note.** The `sfs` subcommand is present in the VariantFlow 1.5.0 release binary. Its folded and unfolded spectra were validated against scikit-allel 1.3.13 and match bin for bin (maximum absolute difference zero); see `benchmark/reports/v30-correctness-crosschecks.md`. An earlier note stating that `sfs` was absent from the prebuilt binary referred to a pre-1.5.0 build and no longer applies.

**What it measures.** The site-frequency spectrum (SFS/AFS): a histogram of how many variant sites have each allele count—the raw signal behind Tajima's D and demographic inference.

```
# Unfolded (derived-allele) spectrum
variantflow sfs "$SNPS" -o chr22.sfs.tsv

# Folded (minor-allele) spectrum, when the ancestral allele is unknown
variantflow sfs "$SNPS" --folded -o chr22.sfs_folded.tsv
```

Use `-folded` when alleles cannot be polarised into ancestral/derived. The unfolded output matches `scikit-allel`'s `allel.sfs`; the folded output matches `allel.sfs_folded`.

#### 6 Interpretation

A cautious reading of what typical values suggest—none is proof on its own.

- **High** $\pi$ means high local diversity. African samples (e.g. YRI) typically show higher $\pi$ than non-African samples, reflecting the older, larger ancestral African population and the out-of-Africa bottleneck.
- **Negative Tajima's D** genome-wide is common in humans and consistent with population expansion; localised strongly negative D can hint at a recent sweep, but confirm independently.
- **Positive Tajima's D** suggests an excess of intermediate-frequency variants (balancing selection or a bottleneck).
- **Fst** between human continental groups is usually modest (often ∼0.1): most human variation is within rather than between populations.

- **LD decaying with distance** is the expected baseline; extended LD can flag low recombination or selection.
- **Missingness near zero** and $F$ **near zero** on this curated panel are healthy sanity checks; large deviations on your own data usually mean a data-quality problem to fix before interpreting anything else.

### 7 Conclusion

Starting from a raw 1000 Genomes chromosome-22 VCF, you downloaded and indexed the data, reduced it to clean biallelic SNPs, and computed a full suite of population-genetics statistics—allele frequency, missingness, heterozygosity, Hardy–Weinberg, nucleotide diversity, Tajima's D, linkage disequilibrium, Fst between two human populations, and (on an all-sites file) the missing-data-aware pixy estimates—with a single tool and standard companions. The results reproduce those of established tools and point toward the classic story of human genetic variation: high diversity, modest between-population differentiation, and signatures of past demographic change. Use them as a starting point for careful, hypothesis-driven analysis rather than as final claims.

[5] Miles A, et al. scikit-allel: Explore and analyse genetic variation. Zenodo software package. https://github.com/cggh/scikit-allel
